# Dynamic Extraction and Tracking of Emitted Cellular Transients resolves low-salience fluorescence events

**DOI:** 10.64898/2026.04.16.718018

**Authors:** Wenli Niu, Yufan Chen, Xia Li, Maïna Garnero, Sambre Mach, Anna Verbe, Margaux Le, Rémi Jousseaume, François David, José-Manuel Cancela, Michael Graupner, Claire Eschbach, Nathalie Rouach, Sabir Jacquir, Micaela Galante, Matthieu Lerasle, Glenn Dallérac

**Affiliations:** Paris-Saclay Institute of Neuroscience, Université Paris-Saclay, CNRS UMR9197, 91400, Saclay, France; Karlsruhe Institute of Technology, Karlsruhe, Germany; Center for Interdisciplinary Research in Biology, Collège de France, CNRS UMR 7241, INSERM U1050; Doctoral School N° 158, Sorbonne Université, Paris, France; Université Paris Cité, CNRS, Saints-Pères Paris Institute for the Neurosciences, F-75006 Paris, France; ENSAE, Institut Polytechnique de Paris, France

## Abstract

Genetically encoded fluorescent sensors have expanded our ability to image cellular activity and transmitter release. Yet, sparse and low-salience events remain difficult to resolve against complex and fluctuating fluorescence backgrounds. Here we introduce DETECT, Dynamic Extraction and Tracking of Emitted Cellular Transients, which combines adaptive background suppression, probabilistic classification and multi-object tracking to extract fluorescence events while preserving their identity. Across synthetic datasets, DETECT improved detection and segmentation accuracy and reduced computational cost relative to established event-based methods. We validated DETECT across confocal, two-photon and miniscope imaging, ex vivo and in vivo, using calcium indicators and monoamine sensors. Beyond its technical performance, DETECT extends event-based analysis to low-salience fluorescence signals while resolving events spanning broad ranges of amplitude, morphology and dynamics. By resolving spontaneous dopamine and noradrenaline signals as distinct, trackable release events, DETECT reveals the spatiotemporal organization of neuromodulatory activity and provides a broadly applicable approach to quantitative fluorescence analysis.

## Introduction

Optical imaging has become a central approach for studying brain function, providing direct access to the spatiotemporal dynamics of intracellular activity and extracellular transmitter release^1,2^. Imaging approaches such as confocal and two-photon microscopy, as well as portable miniscopes, enable recordings in acute slices, head-fixed animals, and freely moving preparations^3–7^. A widely used example is the detection of Ca²⁺ transients with genetically encoded calcium indicators such as GCaMP in neurons and glial cells, both *ex vivo* and *in vivo*^8,9^. More recently, GPCR-based fluorescent sensors have extended this capacity to major neurotransmitters including glutamate and GABA, and to neuromodulators such as monoamines, neuropeptides, acetylcholine, and ATP^10–15^. These tools open the possibility of mapping the spatiotemporal dynamics of local transmitter release and neuronal activity in unprecedented detail, and of relating them directly to behavior^11,16–20^. They also generate large, high-dimensional datasets containing fluorescence events that vary widely in amplitude, morphology and dynamics. Sparse, low-salience events are particularly difficult to resolve against complex and fluctuating fluorescence backgrounds. Such events are biologically important because they report ongoing, spatially distributed activity that is not captured by analyses focused on large or stimulus-locked responses. Yet characterizing their occurrence and spatiotemporal organization is essential for determining how discrete local events contribute to basal neuromodulatory states.

Several factors contribute to this complexity. Motion artefacts, hemodynamic signals, and unspecific fluorescence changes can obscure neural activity, while long recordings are affected by baseline drift arising from fluctuations in illumination, detector sensitivity, or environmental conditions^21–23^. Repeated laser exposure further introduces photobleaching, progressively reducing signal intensity and complicating longitudinal measurements. Beyond these experimental constraints, the datasets themselves are inherently high-dimensional as each frame contains tens of thousands of pixels whose dynamics reflect both biological activity and irrelevant signals. The analytical challenge is therefore not only to separate genuine events from background variability, but also to preserve their identity as they emerge, propagate, disappear and recur.

A range of analysis strategies have been developed to address these challenges. Region of interest (ROI)- based approaches such as Suite2P^24^ or CaImAn^25^ achieve efficiency and scalability but are inherently limited by their assumption of spatially static compartments, which restricts their ability to capture propagating or irregular events. Event-based approaches instead model signals at the pixel level, offering richer spatiotemporal fidelity and the possibility to quantify transient phenomena beyond fixed ROIs. The Activity Quantification and Analysis (AQuA) platform^26^ and its successor AQuA2^27^represent important steps in this direction, providing generalizable tools to quantify complex spatiotemporal signals across modalities. However, resolving low-salience events against structured or unstable backgrounds, while preserving their identity across interruptions and changes in morphology, remains challenging and can entail substantial computational cost. Deep learning approaches have also shown promise for segmentation and event detection^28,29^, but they require extensive annotated training sets and carry high computational costs, while their limited interpretability constrains adoption for exploratory studies.

To address these challenges, we developed a novel data-driven pipeline for automatic fluorescence signal extraction and analysis, grounded in machine learning and image signal processing principles. Our Dynamic Extraction and Tracking of Emitted Cellular Transients (DETECT) pipeline combines adaptive background suppression, pixel-wise probabilistic classification and multi-object tracking to extract fluorescence events while preserving their identity over time. By coupling adaptive signal extraction to explicit tracking, DETECT links events across interruptions and changes in spatial organization, allowing their evolution in space and time to be reconstructed. DETECT is broadly applicable to neurotransmitter and calcium imaging, with particular strength for sparse, low-salience events and recordings with complex or unstable backgrounds. Furthermore, to facilitate use by experimental laboratories, we implemented DETECT in a Python-based graphical user interface (GUI) that provides a user-friendly, resource-efficient environment for data processing.

We validated the versatility and efficacy of DETECT across multiple experimental paradigms, including two-photon, confocal, and one-photon miniscope imaging in both *ex vivo* and *in vivo* preparations. Applications to neuronal and astrocytic calcium imaging, together with monoamine sensing, show that DETECT captures a broad diversity of fluorescence events across preparations and imaging modalities. By resolving low-salience spontaneous dopamine and noradrenaline signals as discrete local events, DETECT enables the spatiotemporal organization of neuromodulatory activity to be examined at the event level. These findings establish DETECT as a broadly applicable approach for analyzing cellular and transmitter dynamics.

## Results

### Structure of the DETECT pipeline

DETECT is designed to extend event-based analysis to low-salience fluorescence signals while remaining applicable to diverse forms of cellular activity and transmitter release. To introduce the pipeline and illustrate its performance under challenging conditions, we selected two prototypical datasets: subtle spontaneous dopamine signals and astrocytic calcium activity superimposed on strong, heterogeneous background fluorescence. The first involves detection of spontaneous dopamine transients in acute prefrontal cortex slices, using *ex vivo* expression of the genetically encoded dopamine sensor GRAB_DA2h_ (Fig. 1a). These events are sparse and occur stochastically in space and time, superimposed on a fluorescent baseline that ranges from low (Fig. 1b; Video 1) to high intensity (Fig. 1c, e; Video 2). The second example involves *in vivo* calcium imaging of hippocampal astrocytes in freely moving animals, achieved with GCaMP6f and one-photon miniature microscopy (Fig. 1f, g; Video 3). These cases (Video 2 and 3) were chosen to represent common challenges in fluorescence imaging, such as low signal-to-noise ratio, dynamic background fluctuations or sparse event detection, and provide a basis for describing the full DETECT analysis pipeline.

**Figure 1.**
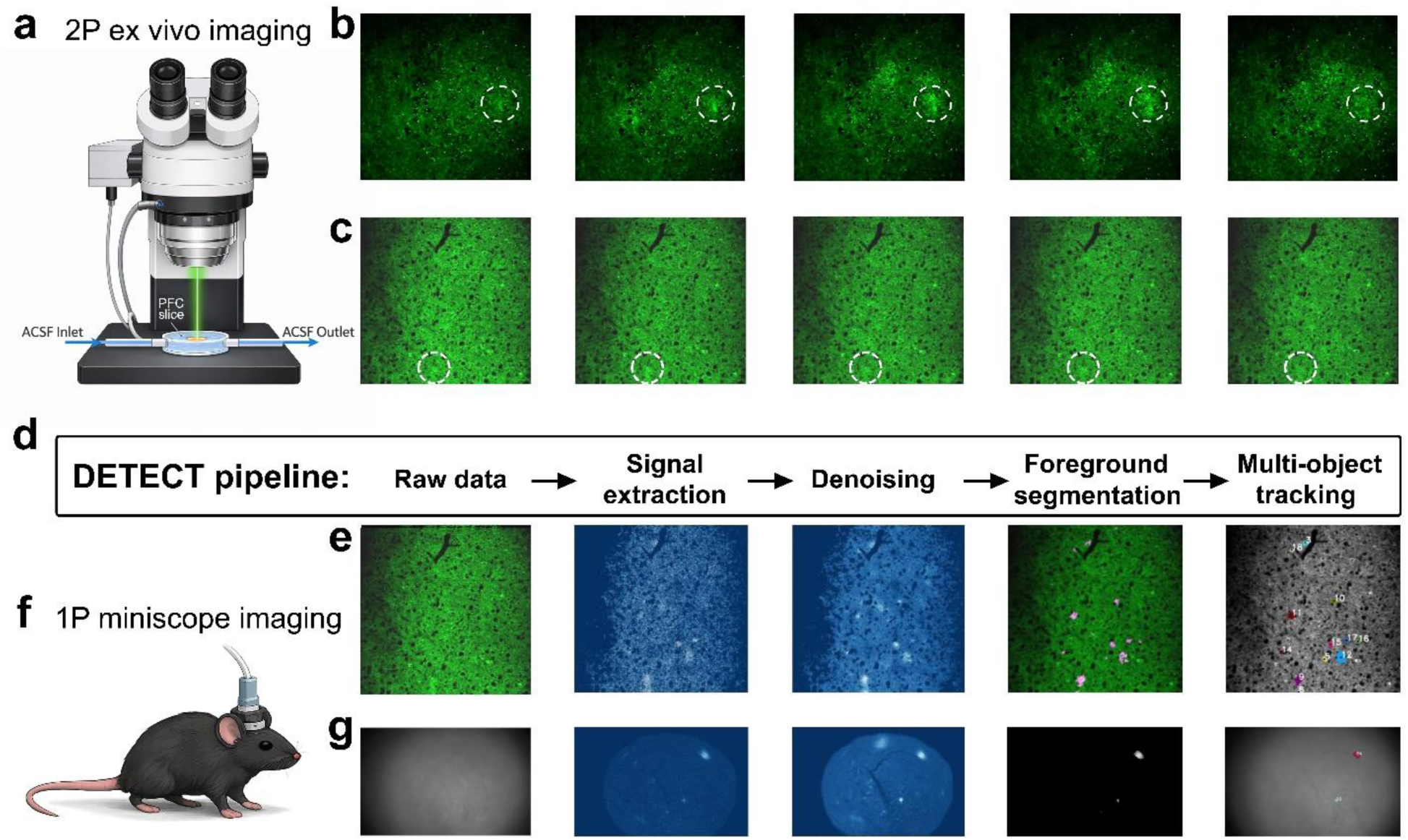
Overview of the DETECT pipeline and representative imaging datasets. **(a)** Schematic of *ex vivo* two-photon imaging in an acute prefrontal cortex slice used for dopamine imaging experiments. ACSF continuously perfuses the recording chamber while fluorescence signals are acquired through the objective. **(b)** Example frames from two-photon imaging of dopamine dynamics in prefrontal cortex slices expressing the GRAB_DA2h_ sensor under low background fluorescence conditions. Transient dopamine release events appear as small diffuse domains emerging from a dim background. Dashed circle indicates a representative event. **(c)** Example frames from two-photon imaging of dopamine dynamics in prefrontal slices expressing the GRAB_DA2h_ sensor under high background fluorescence conditions, illustrating conditions in which dopamine events are barely detectable. Dashed circle indicates a representative event. **(d)** Schematic of the DETECT analysis pipeline showing the sequential processing stages: raw data input, signal extraction, denoising, foreground segmentation, and multi-object tracking. **(e)** Stepwise outputs of the DETECT pipeline applied to a representative two-photon GRAB_DA2h_ dataset, illustrating raw fluorescence data, intermediate signal extraction and denoising stages, foreground segmentation (pink pixels), and final tracking of detected events (colored clusters of pixels). **(f)** Schematic of one-photon miniscope imaging performed in a freely moving mouse. **(g)** Stepwise outputs of the DETECT pipeline applied to a representative miniscope calcium imaging dataset, showing raw fluorescence frames, signal extraction and denoising, foreground segmentation, and tracking of detected Ca²⁺ events.

In order to achieve efficient detection and tracking of dynamics signals in high dimensional datasets, we structured the DETECT pipeline in two main parts: 1/extraction of pixel-level discrete dynamic signals in complex backgrounds and 2/ segmentation and tracking of multiple signals (Fig. 1d). For extraction of relevant signals from background and noise, we used a hierarchical strategy comprising coarse and fine stages. In the coarse stage, we applied Principal Component Analysis (PCA) to categorize signal dynamics by reducing data dimensionality, thereby isolating dominant background and noise components. Unlike conventional PCA approaches that retain principal components, we here subtracted them to reveal the underlying discrete signals. Low-pass filtering is then applied to further remove high-frequency noise. In the finer stages, pixels were classified over time as signal or background by estimating background probability distributions using unsupervised Gaussian mixture models (GMMs), thereby isolating the foreground signals. This machine learning approach enables the distinction of sparse and discrete sensor signals (Fig. 1e, g) from background variations in fluorescence such as photobleaching and laser instability. After successfully extracting dynamic signals, the second part of the DETECT pipeline consists in multi- target segmentation and tracking of the foreground objects. To this end, we use model-based object detection techniques, allowing precise delineation of objects of interest. Finally, to analyze the propagation of dynamic signals, we track each object’s movement path through correlations between consecutive frames using time-series analysis. Overall, DETECT extracts individual events from heterogeneous fluorescence recordings and follows their evolution in space and time.

### Hierarchical extraction isolates dynamic signals from background

#### Coarse stages of background suppression

Fluorescence recordings often display substantial variability in baseline intensity and background dynamics due to differences in sensor expression, illumination, and acquisition settings. In our representative datasets, dopamine transients recorded with GRAB_DA2h_^30^ in acute prefrontal cortex slices vary substantially in amplitude and include subtle events that are weakly separated from the surrounding fluorescence, while one-photon hippocampal astrocyte recordings with GCaMP6f in freely moving mice show strong background fluorescence and spatial inhomogeneity (Fig. 1b, c, g; Video 2-3). To make pixel values comparable across frames and recordings, z-score scaling is first applied, thus normalizing signals relative to their mean and variance, facilitating comparison across datasets with heterogeneous baselines while discarding absolute intensity differences.

Beyond baseline and amplitude variability, a major source of variance arises from structured, low-frequency background fluctuations, masking the discrete and spatially localized events of interest. In both datasets, these background components arise from tissue-wide fluorescence changes, vascular shadows, or laser drift. We address this using PCA to decompose the dataset into its main sources of variance (Fig. 2a). Unlike conventional uses of PCA that retain the dominant components, we subtract the first components that correspond to global background activity. This approach removes the variance-heavy background while preserving the sparse and localized events. As a result, we uncover distinct hotspots in GRAB_DA2h_ slices and clearly delineated astrocytic Ca²⁺ microdomains in miniscope recordings (Fig. 2 b-c).

**Figure 2.**
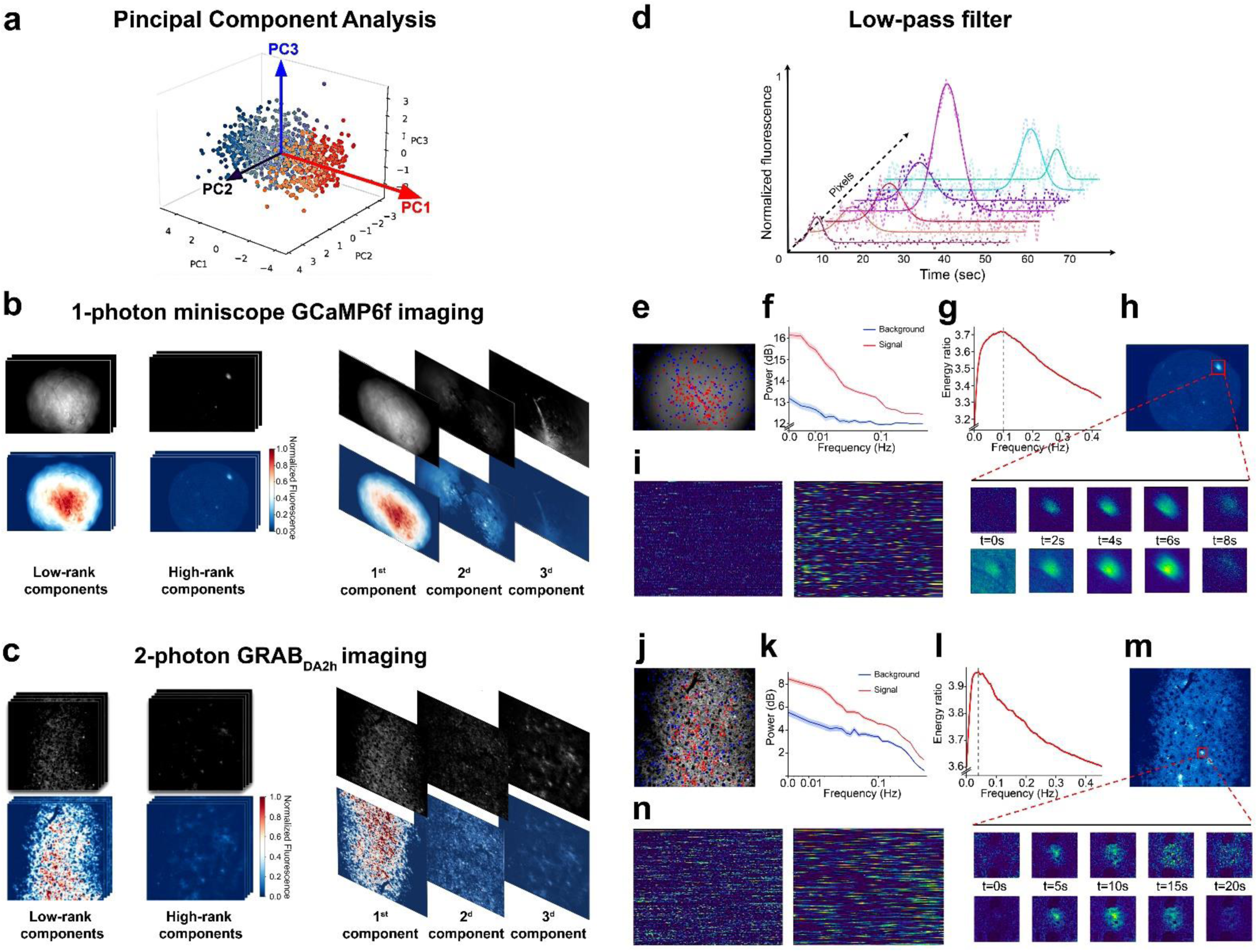
DETECT coarse stages of background suppression. **(a)** Conceptual illustration of principal component analysis (PCA). High-dimensional fluorescence signals are projected onto orthogonal principal components (here PC1–PC3) that capture the dominant sources of variance in the data. **(b)** PCA decomposition of the one-photon miniscope GCaMP6f dataset. Low-rank components mainly capture large-scale background structures and illumination gradients, while high-rank components contain faster fluorescence events corresponding to sparse biological signals in addition to high-frequency noise. The first 3 principal components representing the dominant background contributions are shown and subtracted to isolate discrete transient signals. **(c)** PCA decomposition applied to the two-photon GRAB_DA₂h_ dataset. The first principal components primarily represent the static background contribution as well as slowly varying background structures, and are removed to isolate sparse transient signal events. **(d)** Schematic illustration of low-pass filtering used to suppress high-frequency noise while preserving the temporal dynamics characteristic of biological fluorescence signals. **(e)** Example frame from the miniscope dataset showing pixels classified using convex cone analysis. Pixels identified as signal are shown in red, while randomly selected pixels are shown in blue. One hundred pixels from each category were selected for subsequent spectral analysis. **(f)** Average power spectra of fluorescence fluctuations for background (blue) and signal (red) pixels in the miniscope dataset. **(g)** Frequency-dependent energy ratio derived from the spectral power of signal and background pixels. The peak of the curve identifies the frequency range where signal fluctuations are maximally separated from background noise and is used to determine the cutoff frequency for low-pass filtering (dashed line). **(h)** Representative miniscope fluorescence event illustrating the effect of low-pass filtering. The panel shows the spatial location of the event together with successive frames before filtering (top row) and after filtering (bottom row), demonstrating noise reduction while preserving event structure. **(i)** Pixel-by-time representation of fluorescence dynamics across the miniscope recording before filtering (left) and after filtering (right). **(j)** Example frame from the two-photon dataset showing pixels classified using convex cone analysis. Pixels identified as background are shown in blue and pixels identified as signal are shown in red. One hundred pixels from each category were selected for spectral analysis. **(k)** Average power spectra of fluorescence fluctuations for background (blue) and signal (red) pixels in the two-photon dataset. **(l)** Frequency-dependent energy ratio between signal and background spectra used to determine the optimal cutoff frequency for low-pass filtering. **(m)** Representative two-photon fluorescence event showing the effect of low-pass filtering, with successive frames displayed before (top row) and after filtering (bottom row). **(n)** Pixel-by-time representation of fluorescence dynamics across the two-photon recording before filtering (left) and after filtering (right).

After removal of structured, low-frequency background components, residual high-frequency noise from detector fluctuations or unstable pixels can still obscure the signal. We reduce this by applying a low-pass temporal filter (Fig. 2d), selecting cut-off frequencies from the spectral profiles of representative foreground and background pixels (Fig. 2e-n). This step removes jitter without distorting the onset and decay of events. Together, these coarse-stage operations produce cleaner, more homogeneous datasets in which discrete dynamic events can be reliably isolated in the fine stages of the DETECT pipeline.

#### Fine stages of signal extraction

Following suppression of global background components and removal of high-frequency noise, we refine the detection process to distinguish genuine dynamic events from residual local background fluctuations. As background activity varies unpredictably across both space and time, relying on a single global threshold or fixed probability distribution is inadequate. We therefore implement an adaptive Gaussian mixture model (GMM) that estimates the probability distribution of each pixel’s intensity over the recording. At each time point, the model computes the posterior probability that a given pixel belongs to foreground or background, enabling the identification of brief, spatially restricted events even under unstable illumination or motion- induced artifacts. Importantly, the model parameters, namely component means, variances, and weights, are updated recursively, allowing the algorithm to accommodate gradual drifts, sudden background changes, and the emergence of novel features. Pixels whose estimated background probability falls below an adaptive threshold are classified as foreground, generating a sparse and dynamically updated mask of candidate events (Fig. 3a).

**Figure 3.**
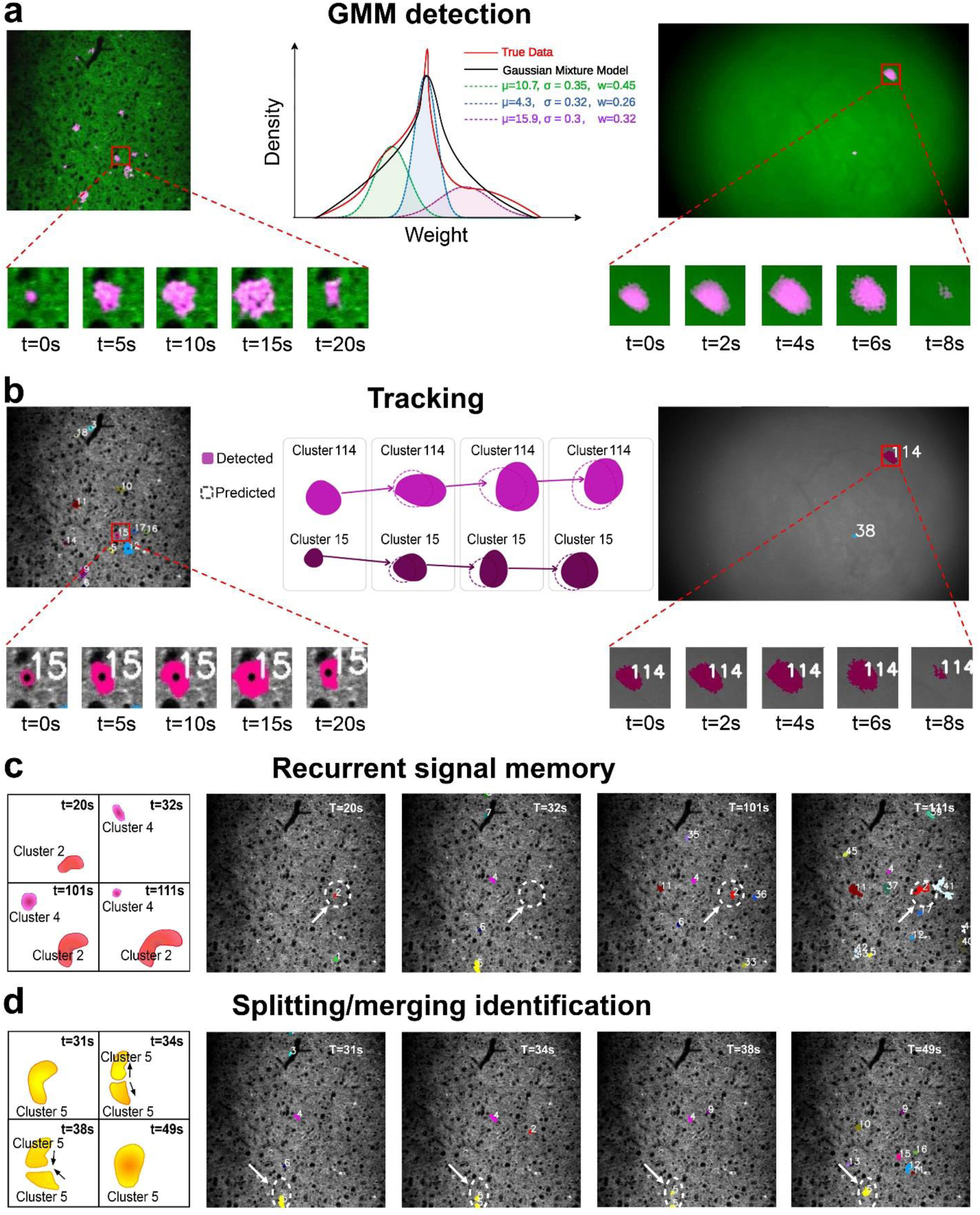
DETECT fine stages of signal extraction. **(a)** Adaptive Gaussian mixture modeling (GMM) classifies pixels as foreground or background based on their temporal intensity distribution. Each pixel is modeled as a mixture of Gaussian components defined by their mean (µ), standard deviation (σ), and mixture weight (w), representing average intensity, variability, and relative contribution to the signal. Pixels classified as foreground (shown in pink) generate a sparse mask of candidate events. **(b)** Adaptive threshold-based segmentation separates spatially adjacent signals while preserving event boundaries. Detected clusters are then associated across frames by matching predicted cluster positions (dashed outlines) with newly detected clusters (solid shapes). **(c)** Optional memory function links recurrent events arising from the same spatial location across time. When a signal temporarily disappears and later reappears within the same region, it is assigned the same cluster identity. **(d)** Identification of splitting and merging events during tracking. When a signal divides into multiple regions or neighboring domains overlap, the resulting regions are associated with the same underlying source identity.

Because fluorescence baselines can drift substantially during long recordings due to photobleaching, illumination fluctuations, or slow biological signals, we avoid classical sliding-window approaches to baseline estimation. Instead, the transition detected by the GMM from background to foreground provides a natural temporal anchor for baseline estimation. For each pixel, the fluorescence value immediately preceding this transition is used as the local reference level, enabling a dynamic ΔF/F calculation that remains robust to baseline instability and does not assume a fixed temporal window.

Applied to our lead datasets, this fine-stage approach reliably isolates discrete signals embedded in complex backgrounds. In GRAB_DA2h_ recordings from acute prefrontal cortex slices, dopamine transients emerge as spatially localized fluorescence clusters that occur stochastically across the field of view (Fig. 3a). In one- photon miniscope recordings of hippocampal astrocytes, localized Ca²⁺ transients are resolved despite superimposed slow baseline waves and spatially inhomogeneous fluorescence (Fig. 3a). This adaptive modeling ensures that only pixels exhibiting statistically distinct temporal dynamics are forwarded to the segmentation and tracking stages, preserving event integrity and minimizing false detections.

### Spatiotemporal segmentation and tracking preserve signal identities

After foreground extraction, the next step is to separate discrete events from one another and to preserve their identity across time. In our representative datasets, dopamine release events in GRAB_DA2h_-labeled prefrontal cortex slices and astrocytic Ca²⁺ domains in miniscope recordings often appear as small, spatially clustered sources surrounded by complex and fluctuating backgrounds. Without robust segmentation and tracking, spatially adjacent sources can be merged, while single sources with intermittent activity can be fragmented into multiple detections. We address this with an adaptive thresholding approach that estimates local thresholds within small neighborhoods, allowing detection sensitivity to adjust to the local intensity context. This prevents bright structures from masking nearby dim events, as in the case of faint astrocytic processes located near strongly active areas. The result is a segmentation that faithfully captures each source’s spatial extent without leakage into neighboring regions (Video 4-5, Fig. 3a). Temporal tracking is then performed with a multi-object association algorithm that links segmented sources across consecutive frames by minimizing the global assignment cost (Video 6-7, Fig. 3b).

An optional memory function extends this process by recognizing when a signal reappears within the same local region after a period of absence and assigning it the same identity (Fig. 3c). This ensures that recurrent events from the same dopamine hotspot or astrocytic microdomain are integrated into a continuous temporal record, preserving the history of each source over the course of an experiment. By maintaining these identities, DETECT allows a distinction between repeated activation of a single signal source and activity arising from distinct sites in close proximity.

In addition, the tracking algorithm explicitly handles topological changes in event morphology. As signals evolve, their spatial footprint may transiently split into multiple regions or merge with neighboring domains. DETECT resolves these configurations during frame-to-frame tracking by computing a centroid-distance cost matrix and identifying optimal assignments using the Hungarian algorithm^31,32^. When several regions appear near the previous centroid of a tracked signal, they are associated with the same source, preserving identity during temporary splitting. Conversely, when regions converge, the merged signal inherits the identity of the closest preceding centroid (Fig. 3d).

These tracking rules ensure that individual signal events remain consistently identified despite interruptions in activity or changes in spatial morphology. In our lead examples, this capability is critical for interpreting patterns of dopamine release or astrocytic Ca²⁺ activity in terms of event-specific dynamics rather than aggregated regional signals. It enables the location, recurrence, and evolution of individual events to be analyzed, providing a foundation for quantifying their synchrony and propagation.

### Additional modules for signal refinement

While the core DETECT pipeline is designed to operate without extensive preprocessing, certain datasets benefit from additional corrections that improve stability and clarity. To accommodate this, we include optional modules for motion correction and spatial filtering that can be activated as needed, depending on the quality of the recordings.

Motion correction addresses the small but cumulative shifts that often occur during imaging. For example, in GRAB_DA2h_ recordings of spontaneous dopamine transients in the prefrontal cortex, slow drift of the field of view can distort spatial alignment, causing recurrent sources to appear at different coordinates over time (e.g., Video 1). DETECT can correct this via a motion correction module using an intensity-based registration approach that maximizes the Enhanced Correlation Coefficient (ECC) between frames. ECC is an iterative algorithm that finds the optimal alignment by maximizing the geometric correlation between pixel intensity patterns, rather than relying on specific landmarks^33^. Unlike feature-based methods that rely on the detection and matching of discrete keypoints, ECC directly operates on the full image intensity distribution, making it particularly well suited for fluorescence imaging data where stable structural features are sparse or absent. In practice, each frame is iteratively aligned to a reference template by estimating an optimal affine transformation that maximizes the correlation between the reference and the transformed frame. This formulation enables subpixel alignment and robust correction of translational, rotational, and mild shear distortions without introducing feature detection bias. This correction is especially critical *for in vivo* recordings, where drift and motion artefacts can introduce apparent displacement of signals across time. By enforcing global intensity consistency, ECC-based correction stabilizes spatial coordinates and preserves the continuity of signal trajectories. As a result, repeated activation of the same source is reliably assigned to consistent locations, preventing misclassification due to mechanical or biological drift (Video 8, Fig. 4).

**Figure 4.**
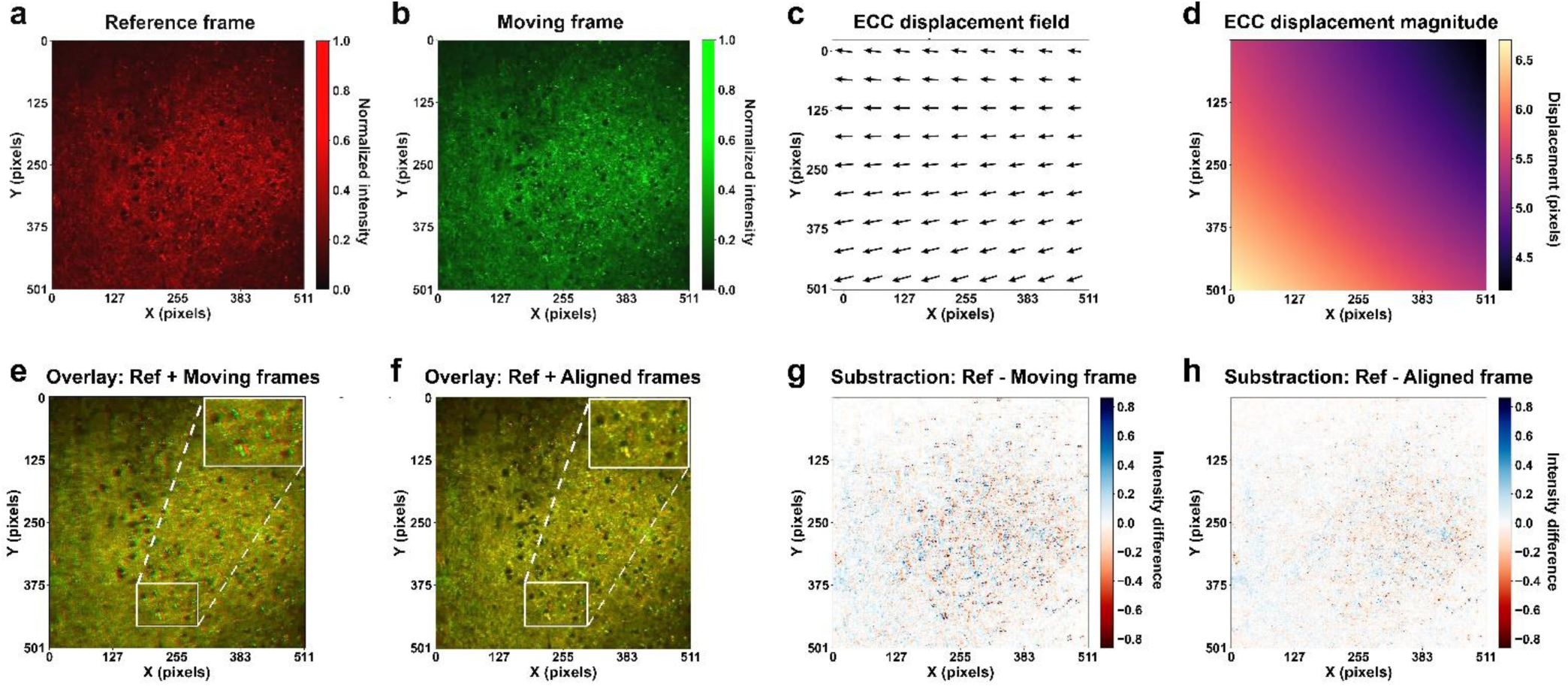
ECC based motion correction. **(a)** Reference frame (frame 0). **(b)** Moving frame (frame 45) prior to alignment. **(c)** Displacement field estimated by the ECC algorithm, shown as a vector field indicating the direction of pixel-wise shifts required to align the moving frame to the reference. **(d)** Corresponding displacement magnitude map, showing the spatial distribution of displacement amplitudes across the field of view. **(e)** Overlay of the reference (red) and moving frame (green) before correction, showing spatial misalignment between corresponding structures, visible as separated red and green signals (inset). **(f)** Overlay of the reference (red) and aligned frame (green) after ECC correction, showing improved spatial correspondence, with yellow regions indicating overlap between red and green signals (inset). **(g)** Pixel-wise difference between reference and moving frame (Ref − Moving), showing widespread positive (red) and negative (blue) intensity differences across the field of view. **(h)** Pixel-wise difference between reference and aligned frame (Ref − Aligned), showing reduced intensity differences and a difference map with minimal residual contrast.

Spatial filtering is also integrated into the DETECT pipeline as an optional processing to improve the visibility of faint structures in datasets with low signal-to-noise ratios. In acute slice recordings of dopamine transients, the fluorescent signal can be partly obscured by detector noise or uneven illumination. Applying moderate Gaussian-based smoothing reduces random pixel fluctuations without blurring the edges of discrete sources, which is essential for accurate segmentation. When combined with the adaptive thresholding in the segmentation step, this produces clearer object boundaries and improves the detection of small, low-intensity events that would otherwise be missed.

Although not required for the functioning of the pipeline, these optional modules extend its applicability to challenging datasets, increasing the robustness of downstream analyses. By stabilizing the field of view and clarifying the spatial profile of sources, they provide a cleaner input for the core detection and tracking stages, ultimately improving the accuracy of event identification and quantification in demanding experimental conditions. This creates an optimal baseline for assessing DETECT’s performance against existing analysis pipelines.

### Performance evaluation of DETECT

#### Comparison with state of the art detection pipelines

To evaluate the performance of DETECT relative to existing event-based analysis pipelines, we compared it with AQuA and AQuA2, the current state-of-the-art event-based pipelines for extracting spatiotemporal fluorescence signals^26,27^. Benchmarking was performed using synthetic datasets with known ground-truth events generated under controlled conditions designed to reproduce the variability and complexity of modern fluorescence imaging recordings (Video 9). Specifically, the synthetic datasets combined spatially nonuniform background fluorescence derived from real imaging data with sparse background transients, onto which agent-based fluorescence events were superimposed. These events varied in amplitude, size-change odds, duration, location-change odds, and propagation, thereby reproducing irregular event morphology and diverse spatiotemporal dynamics observed in experimental recordings.

Detection accuracy was quantified using the mean intersection-over-union (mIoU) between detected and ground-truth events, which measures their spatial and temporal overlap. We also computed the F1 score, i.e. the harmonic mean of precision (the fraction of detected events that are true positives) and recall (the fraction of ground-truth events that were successfully detected), providing a single measure that balances both metrics.

Across the synthetic benchmark datasets, DETECT consistently achieved higher mIoU and F1 scores than both AQuA and AQuA2 (Fig. 5a, b). This advantage was particularly evident in simulations where event morphology evolved dynamically. Increasing the probability of event size changes, which produces expanding or contracting domains across time, led to a marked decline in AQuA2 accuracy, whereas DETECT maintained higher performance across the tested range. Similarly, increasing the number of propagation frames, which generates spatially spreading events, strongly impaired AQuA2 while DETECT preserved substantially better overlap with ground truth. The same ranking was observed when varying event location shifts, with DETECT maintaining the highest mIoU and F1 scores across the tested ranges, followed by AQuA2 and then AQuA (Supplementary Fig. 1a, b). We next examined the effect of background fluctuations, a common challenge in fluorescence imaging. As background amplitude or background fluctuation frequency increased, DETECT again retained higher mIoU and F1 values than both AQuA2 and AQuA across the tested range (Fig. 5c, Supplementary Fig. 1c).

**Figure 5.**
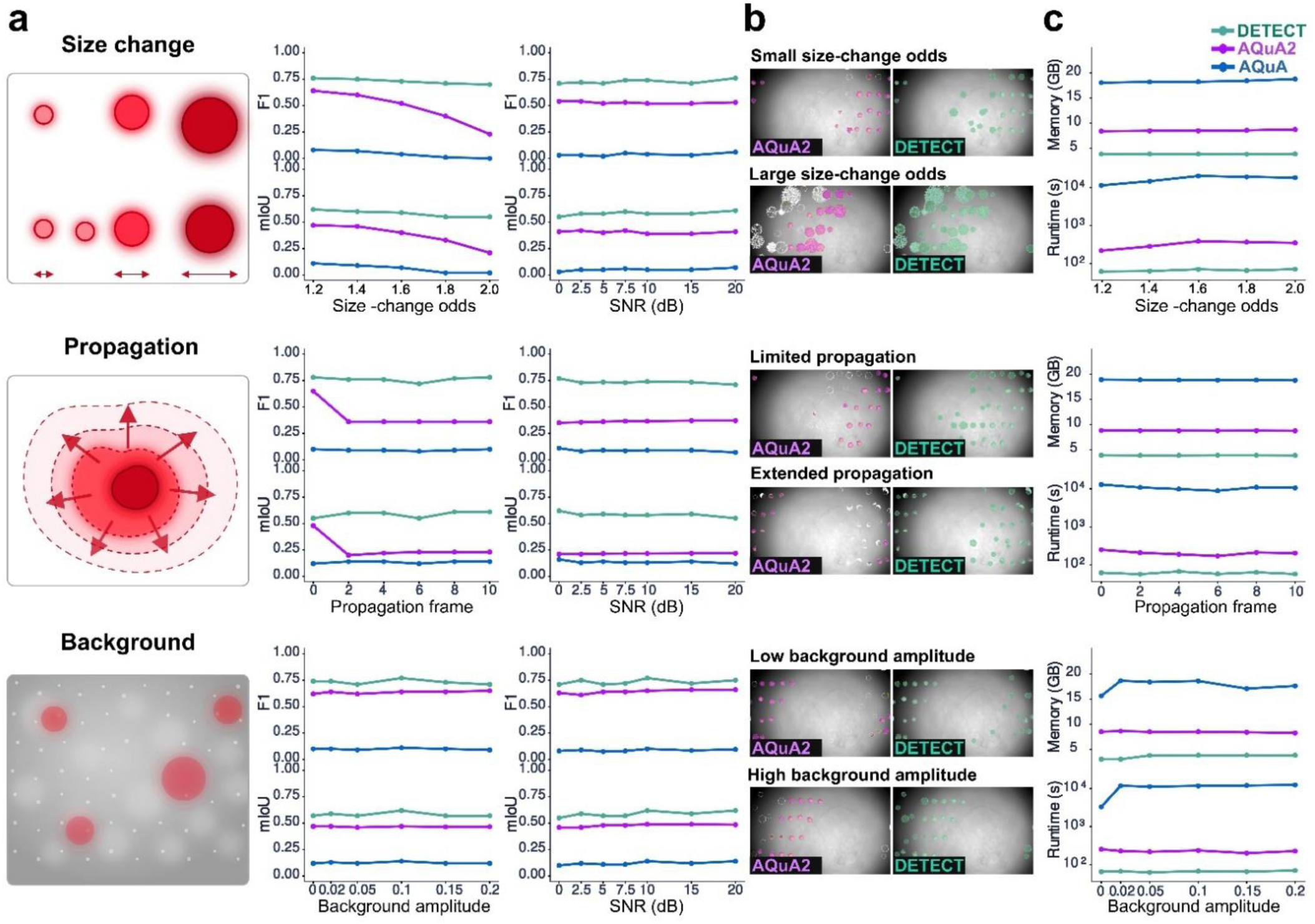
Performance comparison of DETECT with current event-based fluorescence analysis pipelines. **(a)** Performance across simulated conditions. Synthetic datasets were generated with controlled variations in event size change probability, spatial propagation, and background fluctuation amplitude. Detection performance was quantified using F1 score (top) and mean intersection-over-union (mIoU; bottom) across increasing parameter values and signal-to-noise ratios (SNR). DETECT (green) consistently maintained higher detection accuracy than AQuA2 (magenta) and AQuA (blue) across tested conditions. **(b)** Representative detection examples from synthetic datasets. Comparisons between AQuA2 and DETECT under small versus large size-change probabilities, limited versus extended propagation, and low versus high background amplitude. Colored overlays indicate detected events. **(c)** Computational resource usage. Memory consumption (top) and runtime (bottom) measured across simulations with increasing size-change probability, propagation length, and background amplitude. DETECT required substantially less memory and processing time than AQuA2 and AQuA.

Importantly, in addition to improved accuracy, DETECT also displayed substantially lower computational cost. Runtime and memory usage remained low and largely stable across parameter sweeps, whereas AQuA required substantially longer processing times and larger memory allocation. AQuA2 showed intermediate performance but remained consistently slower than DETECT across all simulated conditions.

Overall, these results show that DETECT combines higher segmentation accuracy with substantially improved computational efficiency, providing a robust solution for analyzing complex spatiotemporal fluorescence datasets.

#### Validation across imaging modalities and models

Having established DETECT’s performance on synthetic datasets, we next examined its ability to resolve biologically diverse fluorescence events under experimental conditions. We applied DETECT and AQuA2 to six datasets spanning different sensors, species, brain regions, preparations, and imaging modalities. Because parameter selection can substantially influence event detection, AQuA2 was evaluated using two configurations. The default configuration used the parameter set recommended for the corresponding data type, whereas the adjusted configuration used dataset-specific parameters selected to maximize the recovery of visually identifiable events while limiting detections attributable to background fluctuations. DETECT parameters were likewise adjusted for each dataset using the same criterion. Comparisons across these experimental datasets focused on the resulting event detection and processing time. We first analyzed spontaneous monoamine signals, which combine low event salience with slow or spatially heterogeneous background fluctuations, before extending the comparison to neuronal and astrocytic calcium activity. In two-photon imaging of dopamine dynamics in the mouse prefrontal cortex using GRAB_DA2h_ sensors, the signal consisted of transient dopamine events embedded in slowly varying baseline fluorescence. DETECT identified 88 events in 26.7 s, whereas AQuA2 identified 108 events with adjusted parameters and 317 events with default parameters, with corresponding runtimes of 67.6 s and 643.9 s, respectively (Fig. 6a). Although both methods were sensitive to parameter choice in this dataset, DETECT maintained stable detection across a wider parameter range, whereas AQuA2 configurations either missed a substantial fraction of visually apparent events or produced numerous additional detections not readily attributable to identifiable fluorescence transients (Fig. 6a, Video 10).

**Figure 6.**
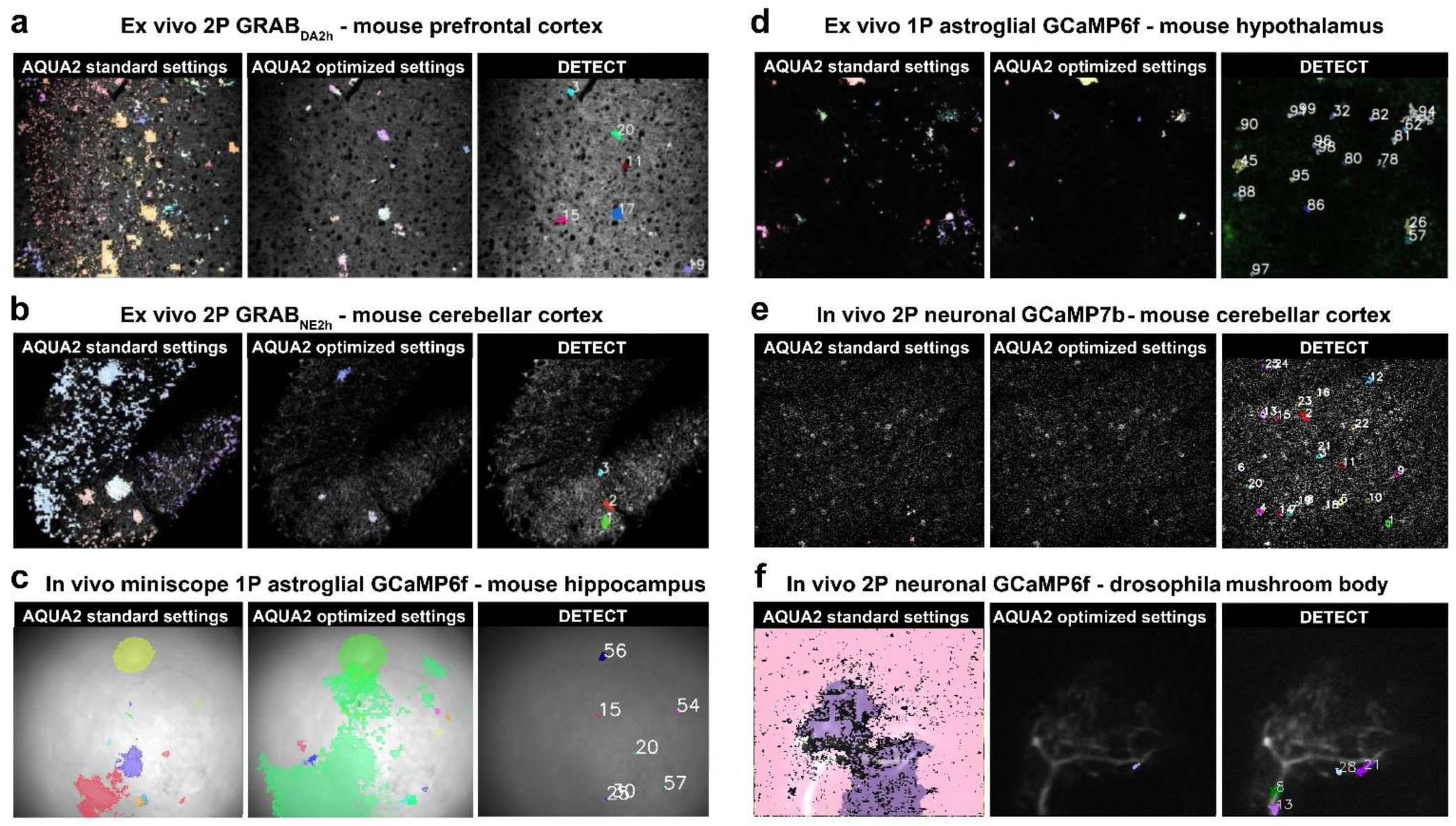
DETECT performance across imaging modalities and models. **(a)** *Ex vivo* two-photon GRAB_DA2h_ imaging in mouse prefrontal cortex. Representative detection outputs obtained with AQuA2 using standard parameters, AQuA2 using optimized parameters, and DETECT. **(b)** *Ex vivo* two-photon GRAB_NE2h_ imaging in mouse cerebellar cortex. Representative detection outputs obtained with AQuA2 using standard parameters, AQuA2 using optimized parameters, and DETECT. **(c)** *In vivo* one-photon miniscope astroglial GCaMP6f imaging in mouse hippocampus. Representative detection outputs obtained with AQuA2 using standard parameters, AQuA2 using optimized parameters, and DETECT. **(d)** *Ex vivo* one-photon astroglial GCaMP6f imaging in mouse hypothalamus. Representative detection outputs obtained with AQuA2 using standard parameters, AQuA2 using optimized parameters, and DETECT. **(e)** *In vivo* two-photon neuronal GCaMP7b imaging in mouse cerebellar cortex. Representative detection outputs obtained with AQuA2 using standard parameters, AQuA2 using optimized parameters, and DETECT. **(f)** *In vivo* two-photon GCaMP6f imaging of a dopaminergic neuron in the *Drosophila* larvae central brain. Representative detection outputs obtained with AQuA2 using standard parameters, AQuA2 using optimized parameters, and DETECT.

In *ex vivo* two-photon GRAB_NE2h_ imaging from mouse cerebellar cortex slices, noradrenaline transients were superimposed on a slowly drifting fluorescence baseline. DETECT identified 52 events in 41.5 s, whereas AQuA2 identified 33 events with adjusted parameters and 155 events with default parameters, with corresponding runtimes of 52.0 s and 99.8 s, respectively (Fig. 6b). Here again, both pipelines efficiency rely on parameter selection, but DETECT preserved detection of genuine events with limited false positives across a broader parameter range, whereas AQuA2 did not yield a parameter configuration that simultaneously captured genuine events while limiting false detections (Video 11).

The same limitation was observed in hippocampal calcium imaging using a head-mounted miniscope, where low signal-to-noise ratio, substantial motion artefacts and strong background fluorescence from out-of-focus planes further complicate detection (Fig. 6c, Video 12). DETECT identified 239 events in 270.52 s, while AQuA2 measured 216 events with adjusted parameters and 94 events with default parameters, processed in 483.871 s with adjusted parameters and 237.958 s with default parameters, and did not provide a parameter configuration that both limited spurious detections arising from background fluctuations and preserved genuine events.

In *ex vivo* confocal recordings of GCaMP6f activity in the mouse hypothalamus, calcium events were sparse but high-amplitude, with low background fluorescence (Video 13). DETECT isolated 182 events in 27.48 s, while AQuA2 measured 284 events with adjusted parameters and 261 events with default parameters, processed in 72.408 s with adjusted parameters and 84.029 s with default parameters. Performance was thus closer between the two pipelines for this dataset (Fig. 6d).

*In vivo* two-photon recordings of molecular layer interneuron activity in the cerebellar cortex displayed discrete somatic calcium transients. DETECT captured 58 events in 196.87 s, while AQuA2 measured 4 events with adjusted parameters in 217.428 s and 304 events with default parameters in 228.617 s. Across parameter configurations, AQuA2 did not reliably capture these somatic events, showing both substantial missed detections and additional detections not spatially associated with visually identifiable somata, whereas DETECT detections were consistently localized to cell bodies (Video 14, Fig. 6e).

Finally, two-photon recordings of *Drosophila* larvae expressing genetically encoded calcium indicators were comparable in spatial dimensions to slice datasets but contained higher event propagation speeds and more frequent overlaps. DETECT measured 89 events in 43 s, while AQuA2 measured 15 events with adjusted parameters in 58.2 s and 66 events with default parameters in 1675.933 s. AQuA2 was likewise limited in resolving rapidly propagating and overlapping events in this dataset, whereas DETECT preserved coherent event structure under these conditions (Video 15, Fig. 6f).

Taken together, these analyses show that the distinction between the pipelines was most evident in GRAB- sensor and miniscope recordings combining low-salience events with complex backgrounds. The outputs of the two pipelines were more comparable in the hypothalamic GCaMP6f dataset, where calcium events were relatively prominent and clearly separated from background fluorescence. More generally, AQuA2 performed well under favorable recording conditions but was more difficult to tune consistently across challenging datasets, whereas DETECT produced robust event-resolved outputs across imaging modalities and required substantially shorter processing times. These findings identify low-salience monoamine signals as a distinctive application of DETECT while demonstrating its broader applicability across calcium indicators, biological preparations, and imaging modalities.

### Spatiotemporal scales of spontaneous dopamine events

We next used DETECT to characterize the spatiotemporal properties of spontaneous dopamine events in GRABDA2h recordings. Basal dopamine transmission is commonly described in terms of tonic extracellular levels, yet current models indicate that this apparent continuity should emerge from highly dynamic local release events shaped by diffusion and uptake. The spatial extent and duration of these events nevertheless remain poorly defined experimentally. By resolving low-salience spontaneous GRAB_DA2h_ signals as individual events, DETECT enabled direct measurement of their duration and spatial extent.

We analyzed 573 events detected across 10 recordings from 8 animals. Spontaneous dopamine events lasted on average 16.68 ± 0.52 s, with individual durations ranging from 5 to 135 s (Fig. 7a,c). Maximum area per event averaged 185.26 ± 10.22 µm², with individual values ranging from 23.00 to 2501.50 µm² (Fig. 7d). To express spatial extent on a linear scale, the maximum area of each event was converted to an equivalent circular radius, defined as the radius of a circle with the same area. This yielded a mean equivalent radius of 6.77 ± 0.15 µm, with individual values ranging from 2.71 to 28.22 µm (Fig. 7b,e). The distributions of both duration and spatial extent were broad, revealing substantial heterogeneity among spontaneous dopamine events, with a subset showing prolonged durations and occupying markedly larger spatial domains than most events (Fig. 7c, d).

**Figure 7.**
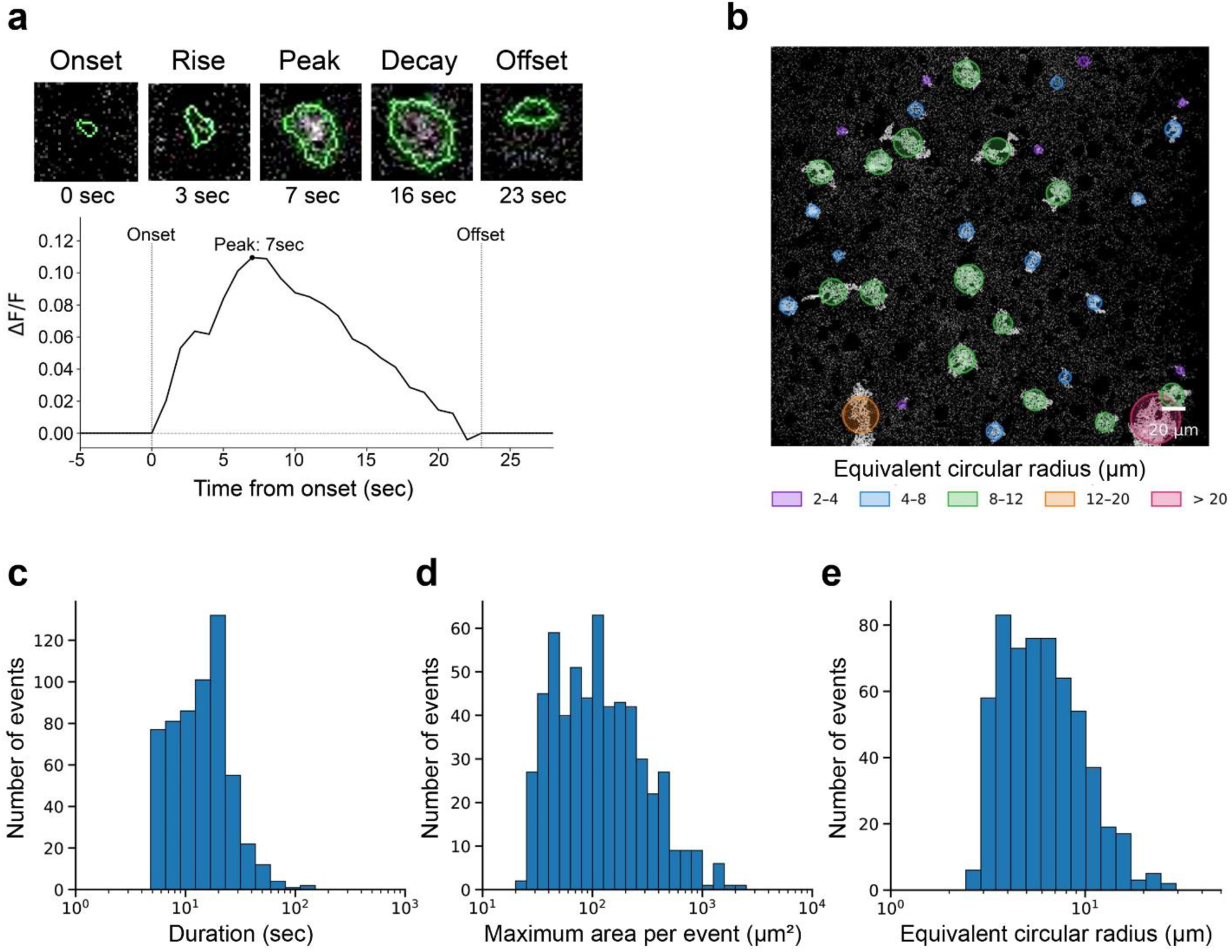
Spatiotemporal properties of spontaneous dopamine events resolved by DETECT. a, Representative spontaneous dopamine event detected in a GRABDA2h recording. Images show the event at onset, during its rise, at peak amplitude, during decay, and at offset. The detected event boundary is shown in green. The corresponding ΔF/F trace is plotted below, with onset, peak, and offset indicated. b, Representative field of view showing the spatial distribution of detected events. Circles are centered on individual events and scaled according to their equivalent circular radius, calculated from the maximum event area. Colors indicate radius ranges of 2-4, 4-8, 8- 12, 12-20, and >20 µm. Scale bar, 20 µm. c, Distribution of event durations across 573 events detected in 10 recordings from 8 animals. d, Distribution of the maximum area reached by each event. e, Distribution of equivalent circular radii calculated from the maximum area of each event. The x axes in c–e are shown on logarithmic scales.

Together, these measurements show that spontaneous dopamine activity comprises heterogeneous events occurring over timescales ranging from seconds to minutes and across a broad range of spatial scales.

DETECT therefore reveals the spatiotemporal properties of spontaneous dopamine signaling, illustrating how its ability to resolve low-salience events can uncover biological features that remain inaccessible to conventional fluorescence analyses and traditional approaches to measuring neuromodulator transmission.

## Discussion

The increasing complexity of fluorescence imaging experiments demands analysis pipelines that operate reliably across modalities, species, and brain regions. Here we introduce DETECT, a broadly applicable and resource-efficient approach for resolving and tracking fluorescence events across diverse imaging conditions, with particular strength for low-salience signals embedded in complex and fluctuating backgrounds. Traditional ROI-based methods are efficient but rely on spatially fixed compartments, limiting their ability to capture events with evolving morphologies or spatial propagation^34^. Event-based approaches such as AQuA and its derivatives^26,27,35^ have established the value of analyzing spatially extended and evolving fluorescence events, but extracting low-salience events and maintaining their continuity across variable backgrounds and changing morphologies remain challenging. DETECT addresses these challenges by integrating PCA-based background suppression with adaptive pixel-wise classification and multi-object tracking. Removing dominant background components at the coarse stage facilitates the extraction of spatially consistent events even in the presence of strong baseline fluctuations. Adaptive classification then separates local events from residual background variability, while tracking preserves event identity across interruptions and changes in spatial organization. Together, these operations enable the location, recurrence, and evolution of individual events to be analyzed across time.

Benchmarking on synthetic datasets that systematically varied event size-change probability, spatial propagation, location shifts, background dynamics, and signal-to-noise ratio showed that DETECT consistently achieved higher segmentation accuracy than AQuA and AQuA2. Performance differences were most pronounced when event morphology evolved across frames, such as during size changes or spatial propagation, conditions under which AQuA and AQuA2 rapidly lost spatial overlap with ground truth whereas DETECT maintained stable mIoU and F1 scores. DETECT also required substantially less runtime and memory across all tested conditions. We focused comparisons on AQuA-based pipelines because AQuA and AQuA2 are the most widely used event-based approaches designed to detect spatially extended fluorescence signals. ROI- and constrained non-negative matrix factorization (CNMF)-based approaches such as Suite2P^24^, CaImAn^25^, and CNMF-E (CNMF for Endoscopic data)^36^ remain highly effective for high-SNR calcium imaging with compact somatic signals, but their underlying assumptions make them ill-suited to low-SNR, spatially diffuse, or morphologically complex events. Event-based methods therefore provide the most relevant benchmark for spatially extended fluorescence signals.

Beyond benchmarking, our analyses demonstrate DETECT’s versatility across diverse experimental paradigms. Across experimental datasets, the distinction between the pipelines was most evident in GRAB- sensor and miniscope recordings combining low-salience events with complex backgrounds. The outputs of the two pipelines were more comparable in the hypothalamic GCaMP6f dataset, where calcium events were relatively prominent and clearly separated from background fluorescence. More generally, AQuA2 produced useful outputs under favorable recording conditions, but in the more challenging datasets we could not identify a parameter regime that recovered visually identifiable events without also generating substantial detections from background fluctuations. In contrast, DETECT produced event-resolved outputs across all tested imaging modalities and consistently required substantially shorter processing times. Together, these findings establish DETECT as a broadly applicable approach for resolving fluorescence events across indicators, biological preparations, and imaging modalities.

Its ability to preserve individual events is especially valuable for spontaneous neuromodulatory signals, whose local spatial and temporal organization remains poorly understood. This is particularly true for monoamines, which signal extensively through volume transmission over scales that are still incompletely defined^37,38^. We examined dopamine as a representative case because existing computational models provide quantitative predictions against which the measured spatial and temporal scales could be compared^39,40^. Tonic dopamine transmission is commonly represented as a slowly varying extracellular baseline, but this description does not resolve the individual events from which that baseline emerges. Microdialysis measures extracellular dopamine over extended tissue volumes and at intervals of several minutes, whereas voltammetric approaches resolve rapid concentration changes only at individual carbon fiber recording sites^41^. Computational studies instead propose that tonic dopamine results from asynchronous release across distributed terminals and the spatial and temporal integration of local concentration transients^42^. However, these models primarily describe average extracellular concentrations or receptor occupancy, leaving the spatial extent and duration of individual spontaneous dopamine events largely inferred rather than experimentally defined.

The measurements obtained here provide an experimental perspective on these scales in the prefrontal cortex. Spontaneous dopamine events typically lasted seconds, with some extending beyond two minutes, and had a mean equivalent radius of 6.8 µm, with the largest event reaching 28.2 µm. Existing models predict concentration domains whose spatial extent depends strongly on the concentration threshold considered. Effective radii of approximately 2 µm and 7-8 µm have been estimated for concentrations associated with low and high affinity dopamine receptor activation, respectively^39^. A more recent model predicted that release of 3000 dopamine molecules produced 1 µM and 100 nM concentration domains extending 1.1 µm and 2.3-2.8 µm, respectively^40^. These predictions were developed for striatal dopamine transmission and should therefore be considered approximate reference points for the PFC, where dopamine clearance relies less on DAT and more substantially on NET, extraneuronal transport, and enzymatic metabolism^43–46^. The mean equivalent radius measured here nevertheless corresponded closely to the broader spatial scale predicted at a low concentration threshold. However, the distribution extended considerably further, reaching 28.2 µm at its upper end and revealing spatial domains beyond the characteristic values derived from isolated release sites.

The temporal comparison revealed a more marked difference. In the computational model, 1 µM transients lasted approximately 2 ms, while 100 nM and 10 nM transients persisted for approximately 8-9 ms and 19- 25 ms, respectively^40^. By contrast, the dopamine signals detected here extended from several seconds to more than two minutes. These durations cannot be attributed solely to the response kinetics of GRAB_DA2h_, whose deactivation occurs on a subsecond timescale. They therefore suggest prolonged or recurrent extracellular dopamine availability, potentially arising from sustained or repeated release, overlapping contributions from multiple sources, and comparatively slow clearance in the PFC.

These comparisons are necessarily approximate. The modeled distances depend on the concentration threshold considered, and the equivalent radius is not a direct measure of molecular diffusion. The detected spatial extent integrates diffusion and clearance, release site geometry, sensor properties, and potentially the simultaneous or sequential recruitment of multiple sources. Similarly, the detected duration reflects the persistence of extracellular dopamine reported by GRAB_DA2h_ rather than the duration of exocytosis at a single release site. The present measurements therefore cannot distinguish among extended diffusion, sustained local release, and overlapping or coordinated contributions from multiple sources.

Nevertheless, the agreement between the mean equivalent radius and the broader scale predicted by existing models suggests that these models capture a characteristic spatial scale of spontaneous dopamine transmission. The broad upper range of the distribution, together with signals persisting for seconds to minutes, reveals additional spatial and temporal heterogeneity that is not captured by isolated, brief transients arising from a single release site. Rather, the observed signals may reflect heterogeneous combinations of diffusion, persistent extracellular dopamine availability, and the recruitment of neighboring sources. To our knowledge, these measurements provide the first event level distributions of both duration and spatial extent for spontaneous dopamine signals across an imaging field. They therefore provide quantitative experimental constraints for models of basal dopamine transmission and begin to bridge the gap between microscopic release models and the slowly varying extracellular levels conventionally described as dopaminergic tone.

More broadly, these findings reveal that spontaneous neuromodulatory activity is structured at the event level. By resolving the location, recurrence, spatial extent, temporal evolution, and coordination of individual signals, DETECT makes it possible to examine how heterogeneous release events combine to generate local dopaminergic tone. This view moves beyond treating tone as a uniform baseline and instead frames it as an emergent property of discrete, spatially extended, seconds long events. DETECT therefore opens a direct route to determining how the organization of spontaneous release shapes neuromodulatory tone across brain regions and behavioral states.

## Methods

### Animals

All experiments were conducted in accordance with the European Community Council Directive 2010/63/EU and approved by the local animal welfare committee (certificate B91272108, Ministère de l’Agriculture et de la Pêche). Male and female C57BL/6J mice aged 8–26 weeks were used. Mice were housed under a 12 h light/dark cycle with *ad libitum* access to food and water.

*Drosophila* experiments were performed in accordance with standard genetic and husbandry procedures approved by the local institutional biosafety committee. Third-instar larvae of both sexes were obtained from established stocks (SS01716-SplitGAL4 > UAS-GCaMP6f) and reared at 25 °C on standard cornmeal medium under a 12 h light/dark cycle.

### Viral injections for GRAB imaging

C57BL/6J mice (P30–P100) were anesthetized with isoflurane and placed in a stereotaxic frame. Adeno-associated viruses encoding genetically encoded monoamine sensors were injected using a glass capillary connected to a Hamilton syringe at controlled rates. The capillary was left in place for 5 min before slow withdrawal to allow diffusion and minimize reflux. For dopamine imaging, AAV9-hSyn-GRAB_DA2h_ (1 µL, 0.2 µL/min) was delivered into the medial prefrontal cortex (AP +2.0 mm, ML ±0.3 mm, DV −2.0 mm relative to bregma). Coronal slices (400 µm) containing the PFC were prepared after 3–4 weeks of expression. For noradrenaline imaging, AAV9-hSyn-GRAB_NE2h_ (500 nL per site, 0.1 µL/min) was injected bilaterally into cerebellar lobule VI (AP −6.8 mm, ML 0 mm, DV −1.4 mm). Parasagittal slices (400 µm) containing lobule VI were obtained 3–4 weeks later for two-photon imaging.

### Ex vivo preparation and imaging

#### GRAB_DA_ imaging in prefrontal cortex slices

Brains were rapidly removed following isoflurane anesthesia and cervical dislocation. Coronal slices (400 µm) containing the medial prefrontal cortex were cut in ice-cold ACSF containing (in mM): 123 NaCl, 2.5 KCl, 1 Na₂HPO₄, 26.2 NaHCO₃, 1.3 MgCl₂, 2.5 CaCl₂, and 10 D-glucose equilibrated with (pH 7.4; 295– 300 mOsm). Slices recovered for 45 min at 30°C in oxygenated ACSF before being transferred to a recording chamber and continuously perfused at 2 ml/min. Imaging was performed on a two-photon microscope (Femtonics) with a 20× water-immersion objective (NA 1.0) at 920 nm, 512 × 502 µm, acquisition rate 30 Hz.

#### GRAB_NE_ imaging in cerebellar slices

Mice aged P60-P130 were deeply anesthetized with isoflurane and sacrificed. The cerebellar vermis was collected after removal of the brain and placed in cold (4°C), oxygenated (95% O_2_ + 5% CO_2_) cutting artificial cerebrospinal fluid (ACSF, in mM: 110 NaCl, 5 KCl, 2.5 CaCl_2_, 1.5 MgSO_4_, 1.24 KH2PO_4_, 10 D-glucose, 27.4 NaHCO_3_, 0.05 D-AP5). The vermis was then parasagitally cut at 400 µm for two-photon imaging. Slices were then transferred to room temperature ACSF for 1h to recover prior to recordings (ACSF, in mM: 110 NaCl, 5 KCl, 2.5 CaCl_2_, 1.5 MgSO_4_, 1.24 KH2PO_4_, 10 D-glucose, 27.4 NaHCO_3_). Imaging was carried out on a two-photon microscope (Femtonics) with a 20× water-immersion objective (NA 1.0) at 920 nm, 512 × 502 µm, acquisition rate 30 Hz.

#### Calcium imaging in hypothalamic astrocytes

GFAP-CreERT2:Ai95 mice (P110 - P140) in which GCaMP6f expression had been induced by tamoxifen (100 mg/kg, i.p., once daily for three consecutive days at P100) were deeply anesthetized with isoflurane and sacrificed. Brains were rapidly removed and placed in cold, oxygenated ACSF containing (in mM): 126 NaCl, 1.5 KCl, 10 HEPES, 21 NaHCO₃, 1.25 MgCl₂, 2 CaCl₂, and 10 D-glucose (pH 7.4; 295–300 mOsm). Coronal slices (300 µm) containing the hypothalamus were cut and transferred to oxygenated ACSF at room temperature for 1 h prior to recordings. Imaging was performed using a confocal microscope (Nikon Eclipse FN1-A1R) with a 40× water-immersion objective (NA 0.8). GCaMP6f was excited at 488 nm and emission collected between 500–550 nm. Images were acquired at 2 Hz with a field of view of 512 × 502 µm.

#### *Drosophila* larvae brain preparation and imaging

Central nervous systems of third-instar *Drosophila* larvae (SS01716-SplitGAL4 > UAS-GCaMP6f) were dissected in a cold buffer containing (in mM): 135 NaCl, 5 KCl, 4 MgCl₂·6H₂O, 2 CaCl₂·2H₂O, 5 TES, and 36 sucrose, pH 7.15 as in^47^, and adhered to poly-L-lysine (SIGMA, P1524)-coated cover glass in small Sylgard (Dow Corning) plates. Imaging was performed in the central brain, in a field centered on a dopaminergic neuron innervating the mushroom body. Two-photon recordings were acquired using a Leica SP8 microscope at 920 nm with a 25× water-immersion objective (NA 0.95). Recordings were acquired at 51 Hz with a field of view of 40.4 × 40.4 µm and a spatial resolution of 0.1579 × 0.1579 µm per pixel.

### In vivo imaging

#### Hippocampal astrocytes imaging

GFAP-CreERT2:Ai95 mice received tamoxifen (100 mg/kg, i.p., three consecutive days at P50) to induce GCaMP6f expression specifically in astrocytes. At P90-P180, a GRIN lens (1×4 mm, ProView™, Inscopix) was implanted above dorsal CA1 (AP −2.0 mm, ML −2.1 mm, DV −1.35 mm, 9° angle). After 2 weeks of recovery and habituation, imaging was performed in freely behaving mice during sleep using a head-mounted miniscope (nVista 3.0, Inscopix) at 15 fps, excitation 0.8–1.2 mW/mm², gain 5-6, field of view 1200 × 800 pixels (1628 × 1018 µm).

#### *In vivo* two-photon calcium imaging of cerebellar molecular layer interneurons

Pvalb-Cre mice (P30–P50; B6.129P2-Pvalbtm1(cre)Arbr/J) were used to enable preferential targeting of cerebellar molecular layer interneurons. Experimental procedures were performed as previously described^48^. Briefly, mice were injected in the left lobulus simplex (AP −6 mm, ML +2 mm relative to bregma) with jGCaMP7b-mRuby3 (ssAAV1/2-hSyn1-dlox-jGCaMP7b-2A-mRuby3(rev)-dlox-WPRE- hGHp(A), diluted 1:5; 3 injections of 300 nL at 200 µm depth). During the same surgical session, a chronic cranial window was implanted over the injection site and a custom stainless steel head post was affixed to the skull. Two to four weeks later, awake mice were habituated to head fixation on a spherical treadmill and underwent *in vivo* two-photon calcium imaging. Recordings consisted of five 60 s trials separated by 2 min intervals during a controlled locomotion paradigm comprising baseline, acceleration, steady locomotion (10 cm/s), and deceleration phases. Two-photon imaging was performed in the molecular layer of the lobulus simplex (20–100 µm below the brain surface) using a custom-built microscope^49^ with a Ti:Sapphire laser (Mai Tai, Spectra Physics) tuned to 910 nm and a 20× water-immersion objective (NA 1.0, Olympus XLUMPlanFl N). Fields of view of 400 × 400 µm were acquired at 30 Hz with a spatial resolution of 0.78 × 0.78 µm per pixel using ScanImage software^50^.

### Simulated dataset

Simulated datasets were generated to benchmark DETECT under controlled conditions. Each synthetic video comprised 260 frames with (800, 1280) pixels and 200 events superimposed on a normalized real background image. Events were generated using an agent-based cluster model rather than a single blob model, with 1200-3500 agents per event, peak amplitudes spanning 35-95% ΔF/F, rapid-rise and exponential-decay temporal waveforms, and event durations of 20-45 frames. Event centers and onset times were distributed across the field of view and recording using a fixed seed for reproducibility, yielding 200 events per video, i.e., 0.77 event onsets per frame on average and up to approximately 42 simultaneously active events within a frame. To increase spatial and temporal heterogeneity, clusters included amplitude jitter, center noise, and per-agent diffusion. Challenge conditions were further introduced by varying cluster size-change odds, location-change odds, and propagation delay. Event propagation was modeled either as cluster translation or as isotropic growth. Background fluctuations were simulated by combining the real background frame with sparse, low-amplitude background perturbations whose amplitudes were expressed relative to the normalized intensity range. Gaussian white noise was added to produce nominal SNRs of 0- 20 dB.

### DETECT pipeline

#### Background suppression

Pixel values were standardized by z-score normalization such that each frame was rescaled to zero mean and unit variance:

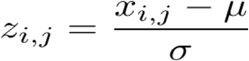

where *x_i, j_* denotes the intensity of pixel *(i, j)*, and where μ and σ denote mean and standard deviation of the frame, respectively.

To isolate background fluorescence, singular value decomposition (SVD) was applied to the centered data matrix:

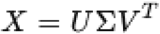

where *X* is the centered data matrix, *U* and *V* are orthogonal matrices containing the left and right singular vectors, respectively, *V^T^* denotes the transpose of *V*, and Σ is the diagonal matrix of singular values. Dominant low-rank components corresponding to background signals were subtracted from the reconstructed dataset.

Residual noise was further reduced by low-pass filtering, with the cut-off frequency estimated using Subspace Convex Cone Analysis (SCCA). Pixels exhibiting high spatiotemporal energy were Fourier-transformed and compared to background pixels to determine the optimal filtering threshold.

#### Adaptive signal extraction

Each pixel was modeled by a Gaussian mixture distribution,

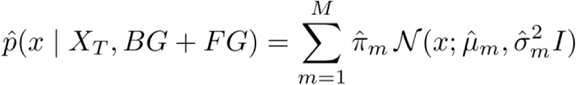

where *x* denotes the pixel intensity value, *M* is the number of Gaussian mixture components, *π̂_m_* is the weight of component *m*, *μ̂_m_* is the corresponding mean, 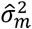 is the variance, and *I* is the identity matrix. The model estimates the probability that each pixel belongs to either background (BG) or foreground (FG) classes over the temporal window X_T_. Mixture weights, means, and variances were recursively updated over time to adapt to baseline drifts, illumination changes, and gradual shifts in fluorescence intensity

#### Segmentation and tracking

Foreground probability maps were segmented using an adaptive Gaussian thresholding method. For each pixel *(x,y)* the local threshold was defined as:

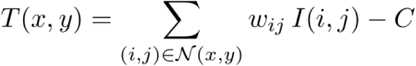

where *N(x,y)* denotes a local neighborhood of size blockSize × blockSize centered on pixel *(x, y)*, *I(i, j)* is the intensity of pixel *(i, j)*, and *C* is a constant used to fine-tune threshold sensitivity.

The Gaussian weighting kernel applied across the neighborhood was defined as:

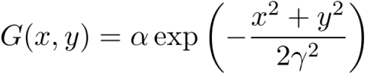

where *α* is a normalization constant and *γ* controls the Gaussian spread. To ensure scale-consistent smoothing across neighborhood sizes, *γ* was defined as a linear function of the neighborhood radius R:

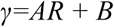

with

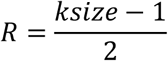

where *ksize* denotes the kernel size, *A* controls the rate at which smoothing increases with neighborhood size, and B sets a minimum level of smoothing independent of neighborhood size (offset). In our implementation, parameters were set to A=0.3 and B=0.8 to provide moderate scaling of the Gaussian spread while maintaining stable baseline smoothing.

Once fluorescence clusters were segmented in each frame, spatiotemporal tracking was achieved using an optimal assignment strategy. For every pair of consecutive frames at *t* and *t+1*, a cost matrix *C_i,j_* was constructed from the Euclidean distances between the geometric centers (centroids) of detected clusters:

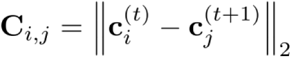

where 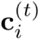 and 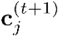 denote the centroid coordinates of clusters in the respective frames. The Hungarian algorithm was then applied to minimize the global cost, yielding the optimal one-to-one assignment of clusters across frames. Signals that failed to match within the predefined temporal or spatial range were classified as new events, whereas clusters that disappeared were marked as terminated. If a cluster reappeared within a set distance of the centroid from its most recent occurrence, it was re-associated with the same signal, thereby allowing continuation across intermittent frames.

Merging and splitting events were handled explicitly. When multiple clusters converged, the merged signal was assigned to the cluster with the closest preceding centroid (“fusion”), while newly merged components were flagged as descendants. Conversely, when one cluster split into multiple subclusters, the offspring were classified as distinct tracks with ancestry preserved. This procedure ensured consistent signal identity across time while accommodating the dynamics of splitting, merging, appearance, and disappearance.

#### Dynamic ΔF/F calculation

For each pixel, the reference fluorescence was defined as the frame immediately preceding the transition timepoint from background to foreground. At event onset, the reference value was set as:

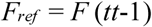

with *tt* the transition timepoint.

The fluorescence change was then computed as:

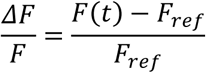

This reference was retained until the pixel returned to background ensuring baselines were anchored locally to each event onset.

## Videos

**Video1. GRAB_DA_ dopamine transients in the prefrontal cortex under low-background conditions.**

DOI: https://doi.org/10.5281/zenodo.19594719

**Video 2. GRAB_DA_ dopamine transients in the prefrontal cortex under high-background conditions.**

DOI: https://doi.org/10.5281/zenodo.19595843

**Video 3. *In vivo* astrocytic calcium imaging with miniscope.**

DOI: https://doi.org/10.5281/zenodo.19596018

**Video 4. Adaptive segmentation of neighboring dopamine transients in the prefrontal cortex.**

DOI: https://doi.org/10.5281/zenodo.19596093

**Video 5. Adaptive segmentation of neighboring calcium signals in miniscope imaging.**

DOI: https://doi.org/10.5281/zenodo.19596200

**Video 6. Multi-object tracking of dopamine transients in prefrontal cortex.**

DOI: https://doi.org/10.5281/zenodo.19596327

**Video 7. Multi-object tracking of calcium signals in miniscope imaging.**

DOI: https://doi.org/10.5281/zenodo.19596485

**Video 8. Motion correction using ECC alignment.**

DOI: https://doi.org/10.5281/zenodo.19596560

**Video 9. Synthetic benchmark dataset for performance evaluation**

DOI: https://doi.org/10.5281/zenodo.19596625

**Video 10. DETECT and AQuA2 comparison in *ex vivo* prefrontal cortex GRAB_DA_ imaging.**

DOI: https://doi.org/10.5281/zenodo.19596752

**Video 11. DETECT and AQuA2 comparison in *ex vivo* cerebellar cortex GRAB_NE_ imaging.**

DOI: https://doi.org/10.5281/zenodo.19596853

**Video 12. DETECT and AQuA2 comparison in miniscope calcium imaging of hippocampal astrocytes.**

DOI: https://doi.org/10.5281/zenodo.19596967

**Video 13. DETECT and AQuA2 comparison in calcium imaging of hypothalamic astrocytes.**

DOI: https://doi.org/10.5281/zenodo.19597075

**Video 14. DETECT and AQuA2 comparison in calcium imaging of cerebellar interneurons.**

DOI: https://doi.org/10.5281/zenodo.19597151

**Video 15. DETECT and AQuA2 comparison in calcium imaging of *Drosophila* larvae central brain dopaminergic neurons.**

DOI: https://doi.org/10.5281/zenodo.19597254

## Video legends

**Video 1. GRAB_DA_ dopamine transients in the prefrontal cortex under low-background conditions.** Two-photon imaging of spontaneous GRAB_DA2h_ fluorescence in acute mouse prefrontal cortex under *ex vivo* baseline conditions. Sparse, spatially restricted dopamine transients appear as discrete and diffuse fluorescence domains emerging from a dim background, illustrating favorable signal-to-noise conditions for event detection. Displayed at 20× real-time speed.

**Video 2. GRAB_DA_ dopamine transients in the prefrontal cortex under high-background conditions.** Two-photon imaging of spontaneous GRAB_DA2h_ fluorescence in acute mouse prefrontal cortex under *ex vivo* baseline conditions. Dopamine events are barely distinguishable from the background, highlighting the need for a robust extraction method. Displayed at 20× real-time speed.

**Video 3. *In vivo* astrocytic calcium imaging with miniscope.** One-photon miniscope recording of GCaMP6f-expressing hippocampal astrocytes in a freely behaving (here sleeping) mouse. Most calcium events are obscured by strong, spatially heterogeneous background fluorescence, and the remaining events are only marginally distinguishable, illustrating the low signal-to-noise conditions of the dataset, hence the need for a robust extraction method. Displayed at 20× real-time speed.

**Video 4. Adaptive segmentation of neighboring dopamine transients in the prefrontal cortex.** Output of the DETECT pipeline after Gaussian mixture model-based foreground extraction and segmentation showing separation of spatially adjacent fluorescence events, with detected foreground pixels highlighted in pink. Local adaptive thresholding preserves event boundaries and prevents merging of nearby sources despite heterogeneous background intensity. Displayed at 20× real-time speed.

**Video 5. Adaptive segmentation of neighboring calcium signals in miniscope imaging.** Output of the DETECT pipeline after Gaussian mixture model-based foreground extraction and segmentation in hippocampal astrocyte miniscope recordings, with detected foreground pixels highlighted in pink. Local adaptive thresholding separates spatially adjacent fluorescence events while preserving event boundaries despite heterogeneous background intensity. Displayed at 20× real-time speed.

**Video 6. Multi-object tracking of dopamine transients in prefrontal cortex.** Tracking of segmented spontaneous dopamine transients in ex vivo two-photon GRABDA2h recordings from mouse prefrontal cortex. DETECT preserves source identity across frames and reconstructs the temporal dynamics of recurrent dopamine hotspots. Displayed at 20× real-time speed.

**Video 7. Multi-object tracking of calcium signals in miniscope imaging.** Tracking of segmented fluorescence events across consecutive frames in hippocampal astrocyte miniscope recordings. Individual sources are assigned consistent identities over time, allowing reconstruction of their temporal dynamics and spatial trajectories. Displayed at 20× real-time speed.

**Video 8. Motion correction using ECC alignment.** Example of motion correction applied to the two-photon recording shown in Video 1, depicting spontaneous dopamine transients in acute mouse prefrontal cortex slices using the GRAB_DA2h_ sensor. Left, original video showing progressive lateral displacement of the field of view over time. Right, the same recording after Enhanced Correlation Coefficient (ECC)-based motion correction, in which frame-to-frame registration stabilizes the imaging field and eliminates apparent displacement across frames. Displayed at 20× real-time speed.

**Video 9. Synthetic benchmark dataset for performance evaluation.** Simulated fluorescence dataset with realistic background fluctuations and ground-truth spatiotemporal events varying in size, propagation, and overlap, used to quantitatively compare the detection accuracy and robustness of DETECT (green), AQuA (blue), and AQuA2 (magenta).

**Video 10. DETECT and AQuA2 comparison in *ex vivo* prefrontal cortex GRAB_DA_ imaging.** Comparison of DETECT and AQuA2 outputs obtained using optimized parameter settings maximizing visually identifiable event detection for each dataset, alongside default AQuA2 parameters recommended for that imaging modality, applied to ex vivo two-photon GRABDA2h imaging in acute mouse prefrontal cortex. Displayed at 20× real-time speed.

**Video 11. DETECT and AQuA2 comparison in *ex vivo* cerebellar cortex GRAB_NE_ imaging.** Comparison of DETECT and AQuA2 outputs obtained using optimized parameter settings maximizing visually identifiable event detection for each dataset, alongside default AQuA2 parameters recommended for that imaging modality, applied to ex vivo two-photon GRABNE imaging in acute mouse cerebellar cortex. Displayed at 20× real-time speed.

**Video 12. DETECT and AQuA2 comparison in miniscope calcium imaging of hippocampal astrocytes.** Comparison of DETECT and AQuA2 outputs obtained using optimized parameter settings maximizing visually identifiable event detection for each dataset, alongside default AQuA2 parameters recommended for that imaging modality, applied to *in vivo* one-photon miniscope recordings of hippocampal astrocytic calcium activity. Displayed at 20× real-time speed.

**Video 13. DETECT and AQuA2 comparison in calcium imaging of hypothalamic astrocytes.** Comparison of DETECT and AQuA2 outputs obtained using optimized parameter settings maximizing visually identifiable event detection for each dataset, alongside default AQuA2 parameters recommended for that imaging modality, applied to ex vivo confocal recordings of GCaMP6f activity in the mouse hypothalamus. Displayed at 20× real-time speed.

**Video 14. DETECT and AQuA2 comparison in calcium imaging of cerebellar interneurons.** Comparison of DETECT and AQuA2 outputs obtained using optimized parameter settings maximizing visually identifiable event detection for each dataset, alongside default AQuA2 parameters recommended for that imaging modality, applied to in vivo two-photon recordings of cerebellar molecular layer interneuron calcium activity monitored using the GCaMP7b sensor. Displayed at real-time speed.

**Video 15. DETECT and AQuA2 comparison in calcium imaging of *Drosophila* larvae central brain dopaminergic neurons.** Comparison of DETECT and AQuA2 outputs obtained using optimized parameter settings maximizing visually identifiable event detection for each dataset, alongside default AQuA2 parameters recommended for that imaging modality, applied to *ex vivo* two-photon recordings of calcium activity in *Drosophila* larvae dopaminergic afferences in the mushroom body, monitored using the GCaMP6f sensor. Displayed at real-time speed.

## Data availability

All datasets generated and analyzed during this study are available from the corresponding authors upon reasonable request. Representative raw imaging videos from each experimental preparation are included. Source data underlying the figures in this manuscript will be provided with the online version of the article.

## Code availability and graphical user interface

The DETECT pipeline and graphical user interface, scripts used for benchmarking and figure generation are available upon reasonable request.

## Acknowledgements

We thank Jérôme Ribot for helpful discussions. This work was funded by Fondation pour la Recherche Médicale (FRM, AJE20181039590), Fondation pour la Recherche sur le Cerveau (FRC, UNAFAM), CNRS, Agence Nationale de la Recherche (ANR-20-CE37-0024, ANR-22-CE16-0009, ANR-22-CE16- 0027-01, ANR-23-CE34-0013-04, ANR-24-CHBS-0007), the Major Research Program of PSL Research University “PSL-Neuro” launched by PSL Research University and implemented by ANR (ANR-10-IDEX- 0001), Institut pour la Recherche en Santé Publique (IReSP, 201134-00), the Sorbonne University doctoral school ED3C and the Chinese Scholarship Council.

## References

1. Markicevic, M., Savvateev, I., Grimm, C. & Zerbi, V. Emerging imaging methods to study whole-brain function in rodent models. Translational Psychiatry 2021 11:1 11, 1–14 (2021).

2. Daria, V. R., Castañares, M. L. & Bachor, H. A. Spatio-temporal parameters for optical probing of neuronal activity. Biophysical Reviews 2021 13:1 13, 13–33 (2021).

3. Cramer, S. W. et al. Through the looking glass: a review of cranial window technology for optical access to the brain. J. Neurosci. Methods 354, 109100 (2021).

4. Ait Ouares, K., Jaafari, N., Kuczewski, N. & Canepari, M. Imaging Native Calcium Currents in Brain Slices. Adv. Exp. Med. Biol. 1131, 73–91 (2020).

5. Cheng, J., McMahon, S. M., Piston, D. W. & Jackson, M. B. Comparing confocal and two-photon Ca2+ imaging of thin low-scattering preparations. Biophysical Reports 3, 100109 (2023).

6. Chen, K., Tian, Z. & Kong, L. Advances of optical miniscopes for in vivo imaging of neural activity in freely moving animals. Front. Neurosci. 16, 994079 (2022).

7. Knöpfel, T. & Song, C. Optical voltage imaging in neurons: moving from technology development to practical tool. Nat. Rev. Neurosci. 10.1038/s41583-019-0231-4 doi:10.1038/s41583-019-0231-4.

8. Tian, L., Akerboom, J., Schreiter, E. R. & Looger, L. L. Neural activity imaging with genetically encoded calcium indicators. Prog. Brain Res. 196, 79–94 (2012).

9. Nasu, Y., Shen, Y., Kramer, L. & Campbell, R. E. Structure- and mechanism-guided design of single fluorescent protein-based biosensors. Nature Chemical Biology 2021 17:5 17, 509–518 (2021).

10. Sun, F. et al. Next-generation GRAB sensors for monitoring dopaminergic activity in vivo. Nat. Methods 17, 1156–1166 (2020).

11. Patriarchi, T. et al. Ultrafast neuronal imaging of dopamine dynamics with designed genetically encoded sensors. Science 360, (2018).

12. Chen, J. et al. The Property-Based Practical Applications and Solutions of Genetically Encoded Acetylcholine and Monoamine Sensors. J. Neurosci. 41, 2318–2328 (2021).

13. Zheng, Y. & Li, Y. Past, Present, and Future of Tools for Dopamine Detection. Neuroscience 525, 13– 25 (2023).

14. Qian, T., Wang, H., Xia, X. & Li, Y. Current and emerging methods for probing neuropeptide transmission. Curr. Opin. Neurobiol. 81, 102751 (2023).

15. Feng, J. et al. Monitoring norepinephrine release in vivo using next-generation GRABNE sensors. Neuron 112, 1930–1942.e6 (2024).

16. Simpson, E. H. et al. Lights, fiber, action! A primer on in vivo fiber photometry. Neuron 112, 718–739 (2024).

17. Sabatini, B. L. & Tian, L. Imaging Neurotransmitter and Neuromodulator Dynamics In Vivo with Genetically Encoded Indicators. Neuron 108, 17–32 (2020).

18. Xia, X. & Li, Y. A high-performance GRAB sensor reveals differences in the dynamics and molecular regulation between neuropeptide and neurotransmitter release. Nature Communications 2025 16:1 16, 1–17 (2025).

19. Wright, E. C., Scott, E. & Tian, L. Applications of functional neurotransmitter release imaging with genetically encoded sensors in psychiatric research. Neuropsychopharmacology 2024 50:1 50, 269–273 (2024).

20. Sun, F. et al. A Genetically Encoded Fluorescent Sensor Enables Rapid and Specific Detection of Dopamine in Flies, Fish, and Mice. Cell 174, 481–496.e19 (2018).

21. Benisty, H., Song, A., Mishne, G. & Charles, A. S. Review of data processing of functional optical microscopy for neuroscience. Neurophotonics 9, 041402 (2022).

22. Bisht, A., Simone, K., Bains, J. S. & Murari, K. Distinguishing motion artifacts during optical fiber-based in-vivo hemodynamics recordings from brain regions of freely moving rodents. Neurophotonics 11, S11511 (2024).

23. Yogesh, B., Heindorf, M., Jordan, R. & Keller, G. B. Quantification of the effect of hemodynamic occlusion in two-photon imaging of mouse cortex. Elife 14, RP104914 (2025).

24. Pachitariu, M., et al. Suite2p: beyond 10,000 neurons with standard two-photon microscopy. bioRxiv 061507 (2017) doi:10.1101/061507.

25. Giovannucci, A. et al. Caiman an open source tool for scalable calcium imaging data analysis. Elife 8, (2019).

26. Wang, Y. et al. Accurate quantification of astrocyte and neurotransmitter fluorescence dynamics for single-cell and population-level physiology. Nat. Neurosci. 22, 1936 (2019).

27. Mi, X. et al. Fast, accurate, and versatile data analysis platform for the quantification of molecular spatiotemporal signals. Cell 188, 2794–2809.e21 (2025).

28. Eom, M. et al. Statistically unbiased prediction enables accurate denoising of voltage imaging data. Nat. Methods 20, 1581–1592 (2023).

29. Islam, M. S., Suryavanshi, P., Baule, S. M., Baek, S. & Glykys, J. A Deep Learning Approach for Neuronal Cell Body Segmentation in Neurons Expressing GCaMP Using a Swin Transformer. eNeuro 10, (2023).

30. Sun, F. et al. Next-generation GRAB sensors for monitoring dopaminergic activity in vivo. Nat. Methods 17, 1156–1166 (2020).

31. Kuhn, H. W. The Hungarian method for the assignment problem. Naval Research Logistics Quarterly 2, 83–97 (1955).

32. Virtanen, P. et al. SciPy 1.0: fundamental algorithms for scientific computing in Python. Nat. Methods 17, 261 (2020).

33. Evangelidis, G. D. & Psarakis, E. Z. Parametric Image Alignment Using Enhanced Correlation Coefficient Maximization. IEEE Trans. Pattern Anal. Mach. Intell. 30, 1858–1865 (2008).

34. Mukamel, E. A., Nimmerjahn, A. & Schnitzer, M. J. Automated analysis of cellular signals from large-scale calcium imaging data. Neuron 63, 747–760 (2009).

35. Wu, Y., Dai, Y., Lefton, K. B., Holy, T. E. & Papouin, T. STARDUST: A pipeline for the unbiased analysis of astrocyte regional calcium dynamics. STAR Protoc. 5, (2024).

36. Zhou, P. et al. Efficient and accurate extraction of in vivo calcium signals from microendoscopic video data. Elife 7, (2018).

37. Özçete, Ö. D., Banerjee, A. & Kaeser, P. S. Mechanisms of neuromodulatory volume transmission. Molecular Psychiatry 2024 29:11 29, 3680–3693 (2024).

38. Liu, C., Goel, P. & Kaeser, P. S. Spatial and temporal scales of dopamine transmission. Nat. Rev. Neurosci. 22, 345 (2021).

39. Rice, M. E. & Cragg, S. J. Dopamine spillover after quantal release: rethinking dopamine transmission in the nigrostriatal pathway. Brain Res. Rev. 58, 303–313 (2008).

40. Wiencke, K., Horstmann, A., Mathar, D., Villringer, A. & Neumann, J. Dopamine release, diffusion and uptake: A computational model for synaptic and volume transmission. PLoS Comput. Biol. 16, e1008410 (2020).

41. Liu, C., Goel, P. & Kaeser, P. S. Spatial and temporal scales of dopamine transmission. Nat. Rev. Neurosci. 22, 345–358 (2021).

42. Dreyer, J. K., Herrik, K. F., Berg, R. W. & Hounsgaard, J. D. Influence of Phasic and Tonic Dopamine Release on Receptor Activation. Journal of Neuroscience 30, 14273–14283 (2010).

43. Käenmäki, M. et al. Quantitative role of COMT in dopamine clearance in the prefrontal cortex of freely moving mice. J. Neurochem. 114, 1745–1755 (2010).

44. Wayment, H. K., Schenk, J. O. & Sorg, B. A. Characterization of Extracellular Dopamine Clearance in the Medial Prefrontal Cortex: Role of Monoamine Uptake and Monoamine Oxidase Inhibition. Journal of Neuroscience 21, 35–44 (2001).

45. Morón, J. A., Brockington, A., Wise, R. A., Rocha, B. A. & Hope, B. T. Dopamine Uptake through the Norepinephrine Transporter in Brain Regions with Low Levels of the Dopamine Transporter: Evidence from Knock-Out Mouse Lines. Journal of Neuroscience 22, 389–395 (2002).

46. Petrelli, F. et al. Dysfunction of homeostatic control of dopamine by astrocytes in the developing prefrontal cortex leads to cognitive impairments. Mol. Psychiatry 25, (2020).

47. Marley, R. & Baines, R. A. Dissection of third-instar Drosophila larvae for electrophysiological recording from neurons. Cold Spring Harb. Protoc. 2011, 1120–1123 (2011).

48. Bao, J. et al. Synergism of type 1 metabotropic and ionotropic glutamate receptors in cerebellar molecular layer interneurons in vivo. Elife 9, 1–22 (2020).

49. Franconville, R., Revet, G., Astorga, G., Schwaller, B. & Llano, I. Somatic calcium level reports integrated spiking activity of cerebellar interneurons in vitro and in vivo. J. Neurophysiol. 106, 1793– 1805 (2011).

50. Pologruto, T. A., Sabatini, B. L. & Svoboda, K. ScanImage: Flexible software for operating laser scanning microscopes. Biomed. Eng. Online 2, 13- (2003).

