## Supplementary figures and images for "Dynamic Extraction and Tracking of Emitted Cellular Transients resolves low-salience fluorescence events"

### Supplementary Figure 1

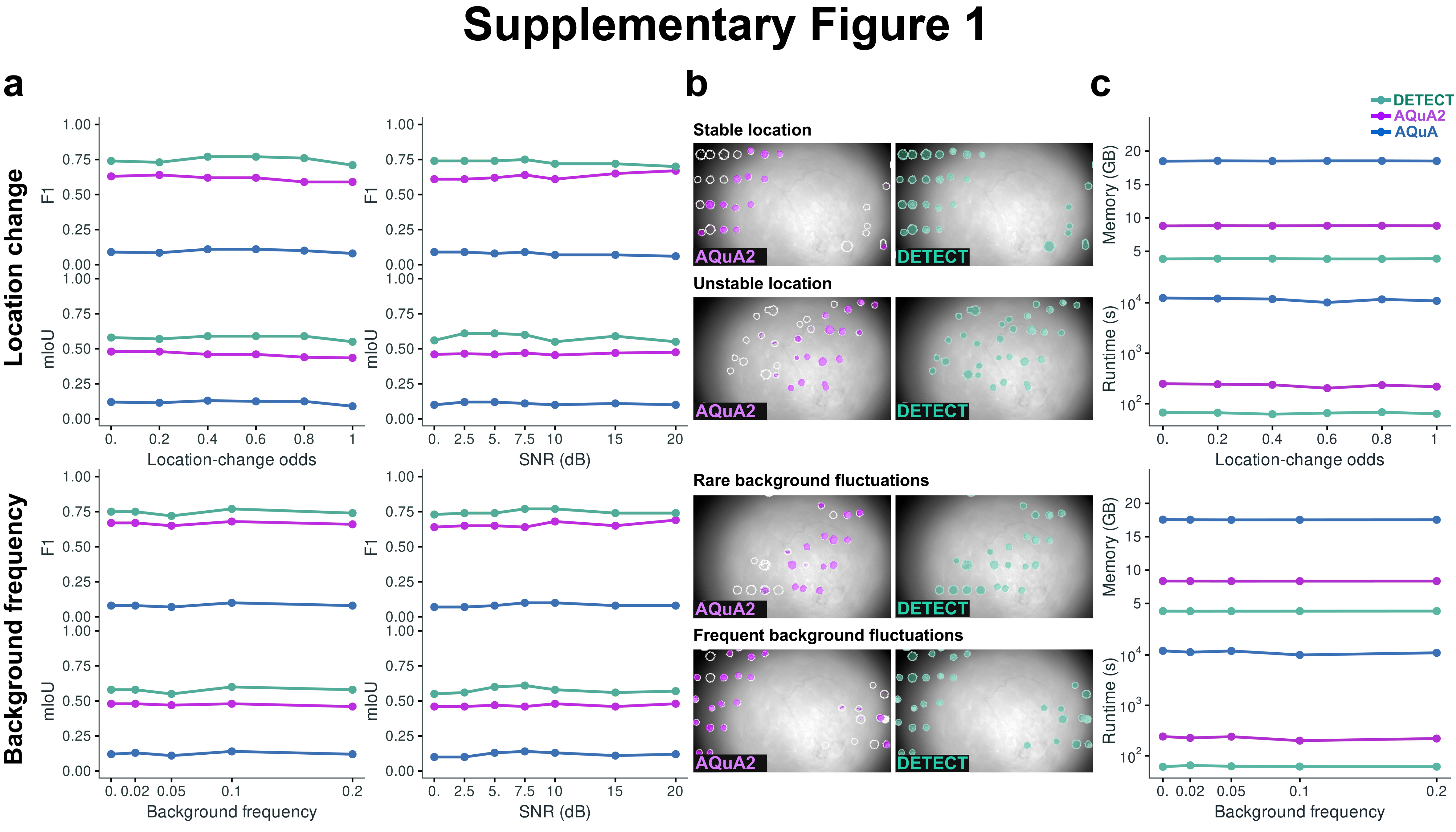
